# Differential effects of the stress peptides PACAP and CRF on sleep architecture in mice

**DOI:** 10.1101/2023.03.22.533872

**Authors:** Allison R. Foilb, Elisa M. Taylor-Yeremeeva, Emma L. Fritsch, Caitlin Ravichandran, Kimberly R. Lezak, Galen Missig, Kenneth M. McCullough, William A. Carlezon

## Abstract

Stress produces profound effects on behavior, including persistent alterations in sleep patterns. Here we examined the effects of two prototypical stress peptides, pituitary adenylate cyclase-activating polypeptide (PACAP) and corticotropin-releasing factor (CRF), on sleep architecture and other translationally-relevant endpoints. Male and female mice were implanted with subcutaneous transmitters enabling continuous measurement of electroencephalography (EEG) and electromyography (EMG), as well as body temperature and locomotor activity, without tethering that restricts free movement, body posture, or head orientation during sleep. At baseline, females spent more time awake (AW) and less time in slow wave sleep (SWS) than males. Mice then received intracerebral infusions of PACAP or CRF at doses producing equivalent increases in anxiety-like behavior. The effects of PACAP on sleep architecture were similar in both sexes and resembled those reported in male mice after chronic stress exposure. Compared to vehicle infusions, PACAP infusions decreased time in AW, increased time in SWS, and increased rapid eye movement sleep (REM) time and bouts on the day following treatment. In addition, PACAP effects on REM time remained detectable a week after treatment. PACAP infusions also reduced body temperature and locomotor activity. Under the same experimental conditions, CRF infusions had minimal effects on sleep architecture in either sex, causing only transient increases in SWS during the dark phase, with no effects on temperature or activity. These findings suggest that PACAP and CRF have fundamentally different effects on sleep-related metrics, and provide new insights into the mechanisms by which stress disrupts sleep.

## INTRODUCTION

Dysfunctional or atypical sleep is part of the diagnostic criteria for stress-related conditions including major depressive disorder (MDD), generalized anxiety disorder (GAD), and post-traumatic stress disorder (PTSD)^1^. Dysfunctional sleep encompasses a range of sleep problems including excessive sleep, diminished sleep, or disrupted and fragmented (i.e., comprising shorter bouts) sleep. Research in humans and laboratory animals have provided extensive evidence that stress and sleep have a reciprocal relationship^2–8^. As an example, stress often dysregulates sleep and circadian patterns ^3–6,9,10^. In turn, abnormal sleep can serve as a form of stress, exacerbating symptom severity in individuals with stress-related conditions^2–5^. Sleep dysregulation in individuals with stress-related conditions are also at increased risk for substance abuse disorders, as many attempt to self-medicate to relieve sleep deficits^11^. An improved understanding of the interactions between stress and sleep will help to advance the ability to diagnose, treat, and even prevent numerous forms of psychiatric illness.

Sleep as a metric has many characteristics, including the ability to measure the same end points across species, that enable enhanced alignment of clinical and neuroscience research on mental health conditions^8,12,13^. Studies in rodents have demonstrated that various forms of stress can produce profound alterations in sleep architecture. For example, shock-based stress or immobilization stress alter diurnal patterns of rapid eye movement sleep (REM) and non-REM sleep^9^. Our lab has examined the effects of chronic social defeat stress (CSDS), an ethological form of stress that involves both physical and emotional elements, on sleep in male mice and found significant alterations in all vigilance states measured: decreases in active wakefulness (AW), increases in slow wave sleep (SWS) and increases in REM^10,14^. In addition, CSDS also disrupted the circadian rhythmicity of body temperature and locomotor activity, such that the normal daily amplitude (rhythm strength) of these endpoints was reduced (flattened)^10^. Importantly, the changes in REM bouts and body temperature persisted beyond the termination of the stressor^10,14,15^. These long-lasting effects of stress are notable because they are commonly observed in individuals with MDD^4,16–19^, and their persistence suggests a high degree of translational relevance in the context of modeling stress-induced psychiatric illnesses, which are by definition persistent and disruptive^1^. The increased use of translationally-relevant endpoints such as sleep and body temperature— among others—in rodents may improve the ability of model systems to more accurately reflect outcomes in humans^8,12,13^.

The mechanisms by which stress triggers psychiatric illness remain unclear^1,8^, which impedes the development of improved therapeutics. Pituitary adenylate cyclase-activating polypeptide (PACAP) and corticotropin-releasing factor (CRF) are peptides with well-validated roles in the biology of stress and stress-related conditions, including mood and anxiety disorders^13,20–25^. Preclinical studies demonstrate that both peptide systems are activated and altered by stress, and administration of either peptide produces stress-like effects^13,20,24^. Both PACAP and CRF are highly conserved across species and produce similar stress-like behavioral effects, including increases in acoustic startle, a measure of vigilance commonly used assess anxiety and fear^21,26–28^. Notably, while acute treatment with CRF produces enhancements in startle and fear responses that resolve within 24 hours of treatment, those produced by acute PACAP treatment can persist for a week or more^21,28–30^. Genetic differences in PACAP and CRF systems are also found in individuals vulnerable to stress^22–24,31,32^. Despite well-characterized sex differences in PACAP and CRF systems that may correspond with the prevalence of stress-related psychiatric disorders in clinical populations, few preclinical studies examining these peptides in parallel have included both males and females^20,22,33–42^. Moreover, the contributions of stress peptides such as PACAP and CRF to stress-induced changes in sleep architecture are not well understood. There is evidence that both PACAP and CRF impact circadian rhythm and sleep, although the findings are often inconsistent or conflicting across studies^9,43–49^. In general, existing studies indicate opposite roles of PACAP and CRF on sleep endpoints such as REM. Some reports indicate that acute CRF treatment reduces REM, whereas PACAP increases REM^43,45–47,50^. The contrasting effects of PACAP and CRF on REM is surprising due to their similar behavioral effects and evidence that PACAP upregulates CRF expression and production^51–53^. Microinfusion of PACAP directly into the pons, a brain region involved in regulation of REM, produces alterations in sleep for a week after treatment, consistent with the persistent effects of PACAP on other behaviors^21,30,47,54^. While stress can produce persistent and often intractable effects on sleep^8^, the ways in which CRF might contribute to these effects have not been thoroughly characterized nor directly compared to those of PACAP^10^.

The goal of the present work was to understand the contributions of PACAP and CRF in the regulation of sleep architecture, body temperature, and locomotor activity in male and female mice. To mimic increases in function occurring during states of stress, we administered doses of each peptide that caused equivalent increases in anxiety-related behavior in a separate assay (the elevated plus maze), thereby enabling physiologically-relevant comparisons. We used a wireless telemetry system and subcutaneous transmitters that enable continuous measurement of electroencephalography (EEG), electromyography (EMG), body temperature, and locomotor activity without tethering that might restrict free movement or posture, or otherwise necessitate abnormal head orientation during sleep^10,14,55,56^. Male and female mice were given a single acute intracerebral ventricular (ICV) infusion of PACAP, CRF, or vehicle (aCSF) and studied for one week after treatment. For each mouse, we used the continuous data sets to quantify vigilance states (AW, SWS and REM), body temperature, and locomotor activity prior to treatment and in 24-hr periods immediately following treatment and one week later. Understanding the long-term impact of PACAP and CRF in male and female mice may identify important differences between the effects of the peptides that could be utilized in the development of new methods to treat or prevent stress-related sleep dysregulation.

## METHODS

### Subjects

Adult (6-8 weeks) male and female C57BL/6 mice were obtained from Jackson Laboratories (Bar Harbor, ME) and were group housed 3-5 per cage until surgery. Colony rooms were temperature-controlled and maintained on a 12-hour light/dark cycle with lights on at 7 am. Mice had ad libitum food and water. All procedures and methods were approved by McLean Hospital Institutional Animal Care and Use Committee and were performed in accordance with the National Institutes of Health’s (NIH) Guide for the Care and Use of Animals.

### Surgery

Mice were anesthetized with intraperitoneal (IP) injections of 100 mg/kg ketamine/10 mg/kg xylazine mixture in saline. Stainless steel guide cannula (26-gauge, P1 Technologies, Roanoke, VA) were implanted for intracerebroventricular (ICV) injection of peptides (relative to bregma: −0.2mm anterior/posterior, +1.0mm medial/lateral, −2.4mm dorsal/ventral). Dummy stylets without projection were used to ensure patency. Transmitters (model F20-EET; Data Sciences International [DSI], St. Paul, MN) were implanted as previously described^10,14,55^. A subcutaneous pocket on the back midway between the neck and hindlegs was made using lubricated forceps and the transmitter was inserted. EMG wires were woven through the trapezius muscle and secured in place with sutures. EEG leads were attached to the skull over the frontal lobe (relative to bregma: +1.00mm anterior/posterior, +1.00mm medial/lateral) and contralateral parietal lobe (relative to bregma: −3.00mm anterior/posterior, −3.00mm medial/lateral) via screws contacting dura. Electrodes and cannula were secured in place using dental cement and the remaining incision was sutured closed. Triple antibiotic ointment was applied to the incision, antibiotic (sulfamethoxazole and trimethoprim) was provided in water, and analgesic ketoprofen (5.0mg/kg) was administered via subcutaneous injection. Mice were single housed at the completion of surgery and given two weeks for recovery before physiological recordings began.

### Treatment

Mice were acclimated to handling a week prior to treatment and stylets were removed and replaced to promote familiarization with the microinfusion procedure. Administration of peptides or vehicle occurred between 9 and 10AM, 2-3 hours after lights-on. Mice were divided into 3 treatment conditions: Vehicle, PACAP, and CRF. Dosages were determined in a separate study that was used to identify concentrations that produced equivalent behavioral (anxiogenic-like) effects in the elevated plus maze (EPM; **see below**). PACAP-38 (0.25μg; Bachem, Torrance, CA) and CRF (1.0μg; Bachem, Torrance, CA) were dissolved in artificial cerebrospinal fluid (aCSF; Harvard Apparatus, Holliston, MA) and administered via an internal cannula projecting 1.0mm beyond the guide cannula at a volume of 1.0μL and a rate of 0.5μL/minute, with an additional 2 minutes to allow for diffusion. Vehicle-treated mice received infusion of aCSF at the same volume and rate as those that received peptide treatment.

### Physiological recordings

Transmitter-implanted mice were housed in standard plastic cages that sat upon receiver platforms (RPC-1; DSI), which allow for wireless data collection in freely moving, untethered animals, as described previously^10,14,55^. Continuous collection of EEG, EMG, locomotor activity and body temperature occurred for four days prior to treatment, and for one week after treatment. Data quantification of vigilance stage—AW, SWS, REM—was determined manually and objectively by a trained scorer blind to treatment condition using Neuroscore (DSI) based on EEG, EMG, and activity recordings.

### Elevated Plus Maze (EPM)

The EPM was used to identify dosages of PACAP and CRF that would produce equivalent behavioral (anxiogenic-like) effects of each peptide, enabling physiologically-relevant comparisons^57^. Vehicle, PACAP, or CRF was administered via ICV cannula as described above, between 9 and 10AM, 30 minutes before testing. As described previously^14^, tests were performed using a standard apparatus (30-cm arms, 5.5 × 5.5-cm center area, 5.5-cm walls, elevated 80 cm); mice were placed in the center to start 5-minute tests. All tests were videotaped and behavior (e.g., time spent on open arms) was quantified using EthoVision (https://www.noldus.com/ethovision-xt).

### Statistical analyses

Data was analyzed using Graphpad Prism 9 with significance set to *P*<0.05 for all analyses. Statistical outliers were identified with ROUT outlier detection test (Q=1); exclusions are noted in Results. As recommended, sexes were combined for the initial analyses, followed by secondary analyses to determine if there were sex differences^58^. EPM behavior was compared across conditions using a one-way ANOVA. Baseline vigilance states were scored as the 24 hours prior to treatment (“Baseline”), and analyses were performed on data expressed as %Baseline during the 24 hours immediately after treatment (10AM-10AM; “Day 1”) and during the same 24-hour window 1 week after treatment (“Day 7”). Changes in vigilance state duration and bouts were compared with mixed effects analyses with repeated measures where applicable. One-way ANOVAs were used to compare changes in EEG absolute power, body temperature, and locomotor activity across treatment conditions. Within each condition and vigilance state, one-sample t-tests were used to compare Day 1 EEG spectral power, as a percentage of Baseline. Significant effects were further analyzed with post-hoc comparisons: Tukey’s post-hoc tests were used to compare across all conditions and timepoints, and where noted, Dunnett’s post-hoc tests were used to make comparisons to Vehicle or Baseline. The EEG data channel was lost in 3 Vehicle-treated male mice before Day 7, preventing vigilance state assessment at that timepoint; however, prior timepoints, as well as all temperature and activity data, is included for these mice.

## RESULTS

### Effects of PACAP and CRF in the EPM

The EPM experiment was designed to identify dosages of PACAP and CRF that produce equivalent anxiogenic-like effects, enabling physiologically-relevant comparisons in the subsequent sleep studies. Initial dosages were selected on the basis of pilot data from studies of these peptides in other procedures (Unpublished data, Carlezon lab). The EPM quantifies anxiety-like states in mice by utilizing innate aversions to bright, open spaces and preferences for dark, enclosed spaces^57^. Mice were placed in the center of the EPM 30 minutes after ICV injection of Vehicle, PACAP (0.25μg), or CRF (1.0μg). ROUT with Q=1 was used to identify outliers, and led to the exclusion of 1 Vehicle male, 2 PACAP males, 1 PACAP female, 1 CRF male, and 1 CRF female; the final numbers of subjects used for the statistical analyses were Vehicle (n=6 males, 6 females), PACAP (n=5 males, 6 females), and CRF (n=7 males, 5 females). Analyses revealed no sex differences—there was no main effect of Sex (F(1,29)=0.44, *P*=0.51, not significant [n.s.]) and no Sex x Condition interaction (F(2,29)=1.11, *P*=0.34, n.s.) (not shown)—providing justification for focusing on the analyses of the data combined across sexes to increase statistical power. With the sexes combined, there was a main effect of Condition (F(2,29)=5.85, *P=*0.007), with both PACAP and CRF reducing the %time on the open arms, relative to vehicle-treated mice (*P*=0.012 and 0.024, respectively; Tukey’s tests) (**Fig. 1**). There were no differences between the PACAP and CRF conditions (*P*=0.95, n.s.)—in fact, the group means were virtually equivalent—providing a justification for selecting these peptide dosages for the subsequent sleep studies.

**Figure 1:**
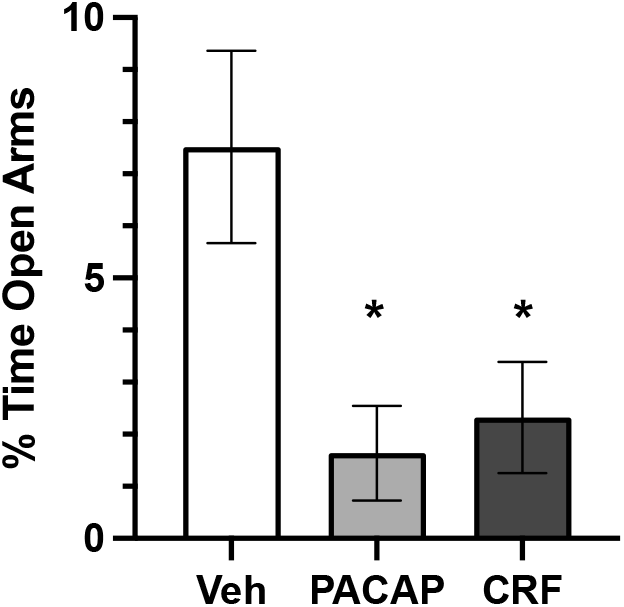
Stress peptide effects on EPM. Percentage of time (±SEM) spent in open arms during 5 minute EPM after treatment with Vehicle (0.0, white), PACAP (0.25 μg, grey) or CRF (1.0 μg, black) in males and females combined. * P < 0.05 compared to Vehicles.

### Baseline Sex Differences in Vigilance States

The overall experimental design is depicted in **Fig. 2A**. First, baseline vigilance states of male and female mice in were assessed in the 24 hours prior to treatment. Unpaired, two-tailed t-tests were used to compare durations of each vigilance state in males (n=27) to those of females (n=25). Analyses indicated significant sex differences in duration of AW (t(50)=2.86, *P*=0.006), and SWS (t(50)=3.22, *P*=0.0009), with increased AW and decreased SWS in females. There were no sex differences in duration of REM (t(50)=1.07, *P*=0.29, n.s.) (**Fig. 2B**).

**Figure 2:**
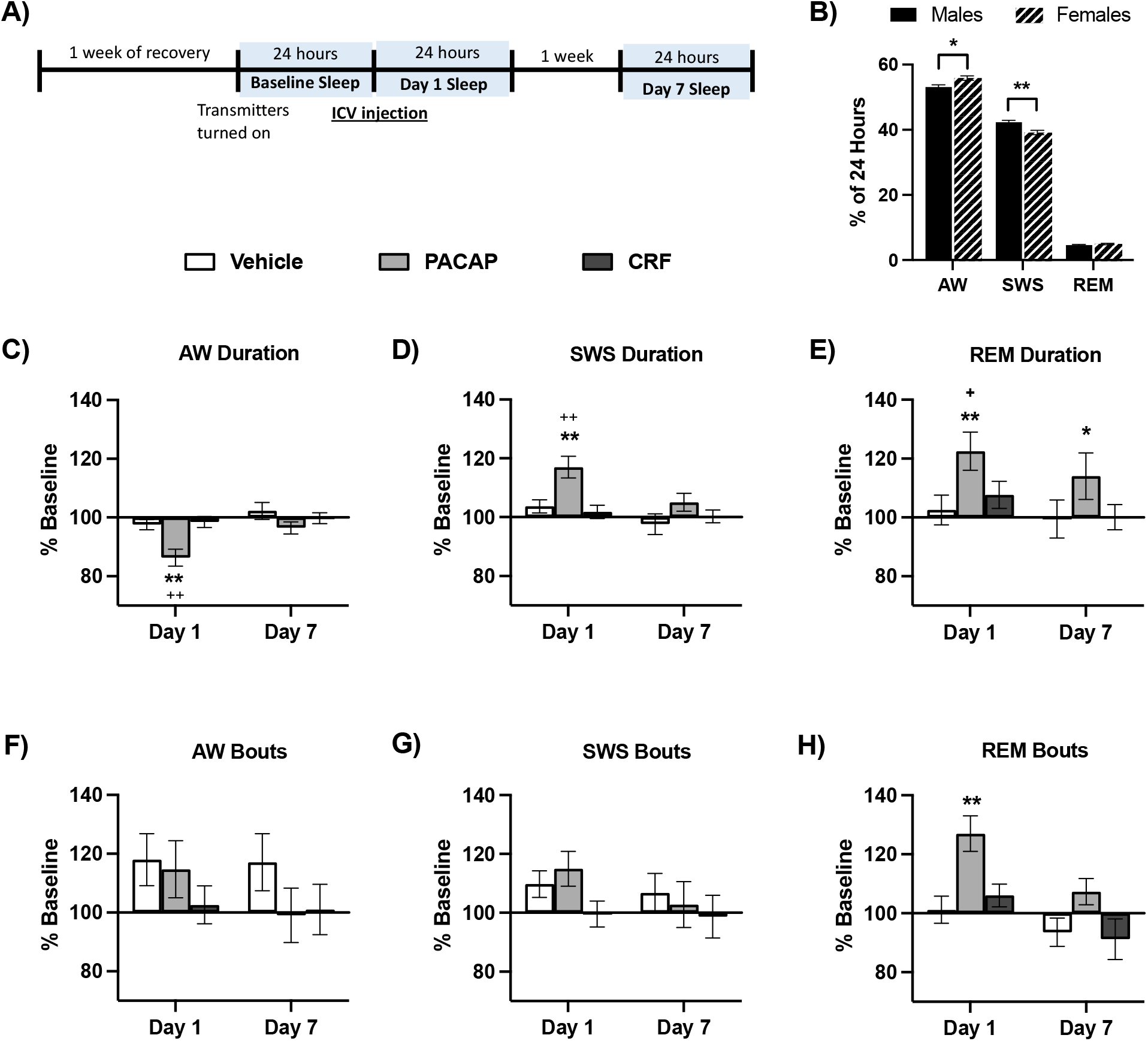
Alterations in vigilance states. **A)** Timeline of experiment and timepoints of vigilance state assessment. **B)** Average (+ SEM) duration of vigilance states at Baseline, prior to peptide treatment with males (solid black bars) and females (striped bars) significantly differing at Baseline in duration of wake (AW) and slow wave sleep (SWS), but not rapid eye movement sleep (REM). **C-H** display changes as percentage of Baseline (± SEM) after Vehicle (white), PACAP (grey) and CRF (black) treatment of **C)** AW duration, **D)** SWS duration, **E)** REM duration, **F)** AW bouts, **G)** SWS bouts, and **H)** REM bouts. Asterisks indicates significant differences from Baseline vigilance states, plus signs indicate significant differences compared to Vehicle-treated animals. * ^+^*P* < 0.05, ** ^++^*P* < 0.01.

### Changes in Vigilance States by Peptide Treatment

As described previously^10^, the effects of stress peptide treatment on vigilance states is calculated as %Baseline. Despite pre-existing sex differences in vigilance state durations, mixed effects analyses of Sex x Timepoint within Condition revealed no sex differences in changes from Baseline after PACAP or CRF treatment. There was a significant main effect of sex on AW and SWS of Vehicle-treated mice; see **Table 1** for all within-Condition Sex x Timepoint comparisons of vigilance state durations. To increase statistical power, males and females were combined in all other analyses to increase the total number of animals per Condition to Vehicle (n=17, comprising 9 males and 8 females), PACAP (n=19, comprising 10 males and 9 females), and CRF (n=16, comprising 8 males and 8 females).

**Table 1:** Mixed effects analyses of Timepoint by Sex within condition and vigilance state. A main effect of sex is found within the Vehicle condition in AW and SWS duration, and a main effect of Timepoint is observed in all vigilance states for PACAP animals. * *P* < 0.05, ** *P* < 0.001

| Condition | Vigilance State | Main Effect of Timepoint | Main Effect of Sex | Timepoint x Sex Interaction |
| --- | --- | --- | --- | --- |
| Vehicle | AW | F (2, 27) = 1.31, $P = 0.29$ | F (1, 15) = 6.55, * $P = 0.022$ | F (2, 27) = 1.75, $P = 0.19$ |
| | SWS | F (2, 27) = 1.51, $P = 0.24$ | F (1, 15) = 5.76, * $P = 0.03$ | F (2, 27) = 1.51, $P = 0.24$ |
| | REM | F (2, 27) = 0.18, $P = 0.83$ | F (1, 15) = 1.02, $P = 0.33$ | F (2, 27) = 0.83, $P = 0.45$ |
| PACAP | AW | F (2, 34) = 12.64, ** $P < 0.0001$ | F (1, 17) = 0.86, $P = 0.37$ | F (2, 34) = 0.43, $P = 0.65$ |
| | SWS | F (2, 34) = 10.17, ** $P = 0.0003$ | F (1, 17) = 0.64, $P = 0.43$ | F (2, 34) = 0.30, $P = 0.74$ |
| | REM | F (2, 34) = 4.62, * $P = 0.017$ | F (1, 17) = 0.02, $P = 0.90$ | F (2, 34) = 0.46, $P = 0.64$ |
| CRF | AW | F (2, 28) = 0.34, $P = 0.71$ | F (1, 14) = 0.84, $P = 0.37$ | F (2, 28) = 0.61, $P = 0.55$ |
| | SWS | F (2, 28) = 0.34, $P = 0.71$ | F (1, 14) = 0.54, $P = 0.47$ | F (2, 28) = 0.72, $P = 0.50$ |
| | REM | F (2, 28) = 2.29, $P = 0.12$ | F (1, 14) = 3.00, $P = 0.11$ | F (2, 28) = 1.96, $P = 0.16$ |

With data from males and females combined, we analyzed durations and number of bouts of each vigilance state by treatment Condition to assess the impact of PACAP and CRF on sleep dysregulation. Changes in duration of sleep reflect excessive or diminished sleep, and increases in SWS or REM bouts represent disrupted or fragmented sleep, as often observed in individuals with stress-related disorders^16,59,60^. A mixed effects analysis of Condition x Timepoint for AW revealed main effects of Timepoint (F(2,95)=10.52, *P*<0.0001), Condition (F(2, 49)=7.45, *P*=0.0015), and a significant Timepoint x Condition interaction (F(4, 95)=4.31, *P*=0.003 (**Fig. 2C**). Post-hoc tests (Tukey’s) indicated that the Day 1 timepoint significantly differed from Baseline and Day 7 (*P*=0.0002 and 0.001, respectively). PACAP-treated animals significantly differed from both Vehicle- and CRF-treated animals overall (*P*=0.0034 and 0.008, respectively) and specifically at Day 1, corresponding with the 24-hour period after treatment (P’s<0.0001). Within conditions, there were no differences between timepoints in Vehicle or CRF-treated animals, but PACAP-treated animals had significant reductions in AW duration at Day 1 compared to Baseline and Day 7 (P’s<0.0001). A mixed effects analysis of AW bouts revealed no significant main effect of Timepoint (F(2,95)=2.13, *P*=0.125 n.s.), Condition (F(2, 49)=1.28, *P*=0.288, n.s.), nor a Timepoint x Condition interaction (F(4,95)=1.35, *P*=0.258, n.s) (**Fig. 2F**). Reductions in duration without significant changes in the number of bouts indicates that the average length of bouts is shorter, reflecting fragmentation.

A mixed effects analysis of SWS duration also found main effects of Timepoint (F(2,95)=9.69, *P*=0.0001), Condition (F(2,49)=7.40, *P*=0.0016), and a significant Timepoint x Condition interaction (F(4,95)=3.74, *P*=0.0071) (**Fig. 2D**). Post-hoc tests (Tukey’s) revealed that SWS at Day 1 significantly differed from Baseline and Day 7 (*P*=0.0003 and 0.002, respectively). PACAP-treated animals significantly differed from both Vehicle- and CRF-treated animals overall (*P*=0.0046 and 0.0062, respectively) and specifically at Day 1 (*P*=0.0002 and <0.0001). Tukey’s tests for effects within conditions, found no differences between timepoints in Vehicle or CRF-treated animals, but PACAP-treated animals had significantly different SWS duration at Day 1 compared to their Baseline (*P*<0.0001) and Day 7 (*P*=0.005). A mixed-effects analysis of SWS bouts revealed no significant effect of Timepoint (F(2,95)=2.51, *P*=0.087, n.s.), Condition (F(2,49)=1.07, *P*=0.35, n.s.), nor a Timepoint x Condition interaction (F(4,95)=0.92, *P*=0.46, n.s.) (**Fig. 2G**). The combination of longer SWS bouts and increased SWS duration suggests a lack of fragmentation.

Analysis of REM duration revealed a main effect of Timepoint (F (2,95)=5.49, *P*=0.006), but no significant effect of Condition (F(2,49)=3.04, *P*=0.057, n.s.), nor a significant Timepoint x Condition interaction (F(4,95)=1.68, *P*=0.16, n.s.) (**Fig. 2E**). Within Conditions there were no significant changes from Baseline in Vehicle- or CRF-treated animals, but significantly different REM durations at Day 1 (*P*=0.004) and Day 7 *(P=*0.039) compared to Baseline in PACAP-treated animals. Mixed effects analysis of REM bouts revealed main effects of Timepoint (F(2,95)=10.56, *P*<0.0001) and Condition (F(2,49)=6.19, *P*=0.004), and a significant Timepoint x Condition interaction (F(4,95)=4.00, *P*=0.005 (**Fig. 2H**). Post hoc comparisons (Tukey’s) revealed that Day 1 significantly differed from Baseline and Day 7 (*P*=0.0012 and 0.0002, respectively). PACAP-treated animals significantly differed from both Vehicle and CRF conditions (*P*=0.009 and 0.015), specifically at Day 1 (*P*’s<0.0001 and *P*=0.0006), as well as compared to CRF at the Day 7 timepoint (*P*=0.046). Within conditions, REM bouts did not change across timepoints for Vehicle or CRF-treated animals, while PACAP-treated animals had significantly increased bouts of REM compared to Baseline at both Day 1 (*P*<0.0001) and Day 7 (*P*=0.0003). Unlike the case with AW, however, the change (increase) in REM bouts corresponds with the increase in REM duration. This suggests consistency in the length of REM bouts (i.e., lack of fragmentation), and that PACAP-treated animals went into REM more frequently at Day 1 and Day 7 compared to Baseline.

### Vigilance State Durations during the Light Cycle

In all mice, the ICV infusions were performed during the third hour of lights-on in a 12-hour light/dark cycle. To better understand when the treatments produced effects on vigilance, we examined behavior during the light and dark phase of the diurnal cycle. Changes during the light phase occurred in the 9 hours immediately after treatment, while the dark phase includes a period of equivalent length (9 hours) during the dark phase. Mixed effects analysis of AW duration in the light phase revealed a significant Timepoint x Condition interaction (F(4,95)=3.77, *P*=0.0069), but no main effects of Timepoint (F(2,95)=2.036, *P*=0.14, n.s.) nor Condition (F(2,49)=2.99, *P*=0.059, n.s.). Post hoc comparisons (Tukey’s) revealed that PACAP-treated animals significantly differed from Vehicle and CRF-treated animals on Day 1 (*P*’s=0.030 and *P*<0.0001, respectively), and that CRF-treated animals displayed significantly increased AW durations on Day 1 when compared to Day 7 (*P*=0.017). During the dark phase of the light cycle, there was a main effect of Timepoint for AW duration (F (2,95)=224.32, *P*<0.0001), with Day 1 significantly decreased compared to Baseline and Day 7 (*P*’s<0.0001).

Analysis of SWS duration in the light phase also revealed a significant Timepoint x Condition interaction (F(4,95)=3.69, *P*=0.0077), but no main effects of Timepoint (F(2,95)=0.39, *P*=0.68, n.s.) nor Condition (F(2,49)=3.16, *P*=0.051, n.s.). Post hoc comparisons (Tukey’s) revealed that PACAP-treated mice differed from Vehicle and CRF-treated animals at Day 1 (*P*=0.039 and 0.0008, respectively). Within conditions, there were no differences between timepoints in Vehicle animals, but PACAP-treated animals had significantly increased SWS during the light phase at Day 1 compared to Baseline (*P*=0.037) and CRF-treated animals had increased SWS at Day 7 compared to Day 1 (*P*=0.013). During the dark phase of the light cycle, there were main effects of Timepoint, (F(2,95)=24.86, *P*<0.0001) and Condition (F(2, 49)=4.34, *P*=0.018) for SWS duration, but no Timepoint x Condition interaction (F(4, 95)=1.77, *P*=0.14, n.s.). Dark-phase SWS significantly differed at Day 1 compared to both Baseline and Day 7 (*P*’s<0.0001). Between conditions, CRF-treated animals significantly differed from Vehicle-treated animals (*P*=0.021).

Analysis of REM duration in the light phase revealed a main effect of Timepoint (F(2,95)=5.50, *P*=0.0055) but not Condition (F(2,49)=0.89, *P*=0.42, n.s.), nor a Timepoint x Condition interaction (F(2,95)=0.51, *P*=0.73, n.s.). Light phase REM duration on Day 7 timepoint significantly differed from Baseline and Day 1 (*P*=0.025 and 0.0077, respectively). During the dark phase, there was again a main effect of Timepoint (F(2,95)=19.11, *P*<0.0001), but not Condition (F(2,49)=1.05, *P*=0.36, n.s.). nor a Timepoint x Condition interaction (F(4,95)=1.27, *P*=0.29. Dark phase REM duration at Day 1 significantly differed from Baseline and Day 7 (P’s<0.0001).

### Body Temperature and Locomotor Activity

In general, changes in body temperature and activity often correspond with changes in vigilance states. Considering that there were minimal changes in vigilance states at Day 7 in any of the conditions, analyses of temperature and activity focused on changes at Day 1, the 24 hours immediately after treatment, compared to Baseline.

Temperature at Day 1 significantly differed across treatment conditions. A One-way ANOVA of Day 1 body temperatures, as a percentage of Baseline revealed a significant effect of Condition, F (2, 49) = 3.39, *P* = 0.042 (**Fig. 3A**). Post hoc comparisons (Tukey’s) did not further identify differences between groups, but Dunnett’s multiple comparisons to directly compare Vehicle treatment to each peptide treatment revealed a significant difference between Vehicle and PACAP-treated animals (*P*=0.049), with no difference between Vehicle and CRF-treated animals (*P*=0.99, n.s.). Within-condition t-tests comparing Baseline to Day 1 revealed no significant changes in Vehicle-treated (t(16)=0.19, *P* = 0.85, n.s.) nor CRF-treated mice (t(15)=0.036, *P*=0.97, n.s.), although there was a significant reduction in temperature in PACAP-treated mice (t(18)=2.33, *P*=0.032). More detailed (1-hr bins) analysis of body temperature revealed characteristic decreases in body temperature during the light phase and increases in temperature at the start of the dark phase, with a dip in temperature midway through the dark phase^10,55,56^. In the hourly temperature data from the PACAP-treated animals, reductions in body temperature occurred during the dark phase of the light cycle, corresponding with treatment-induced increases in sleep when the mice are typically awake and active (**Fig. 4B**).

**Figure 3:**
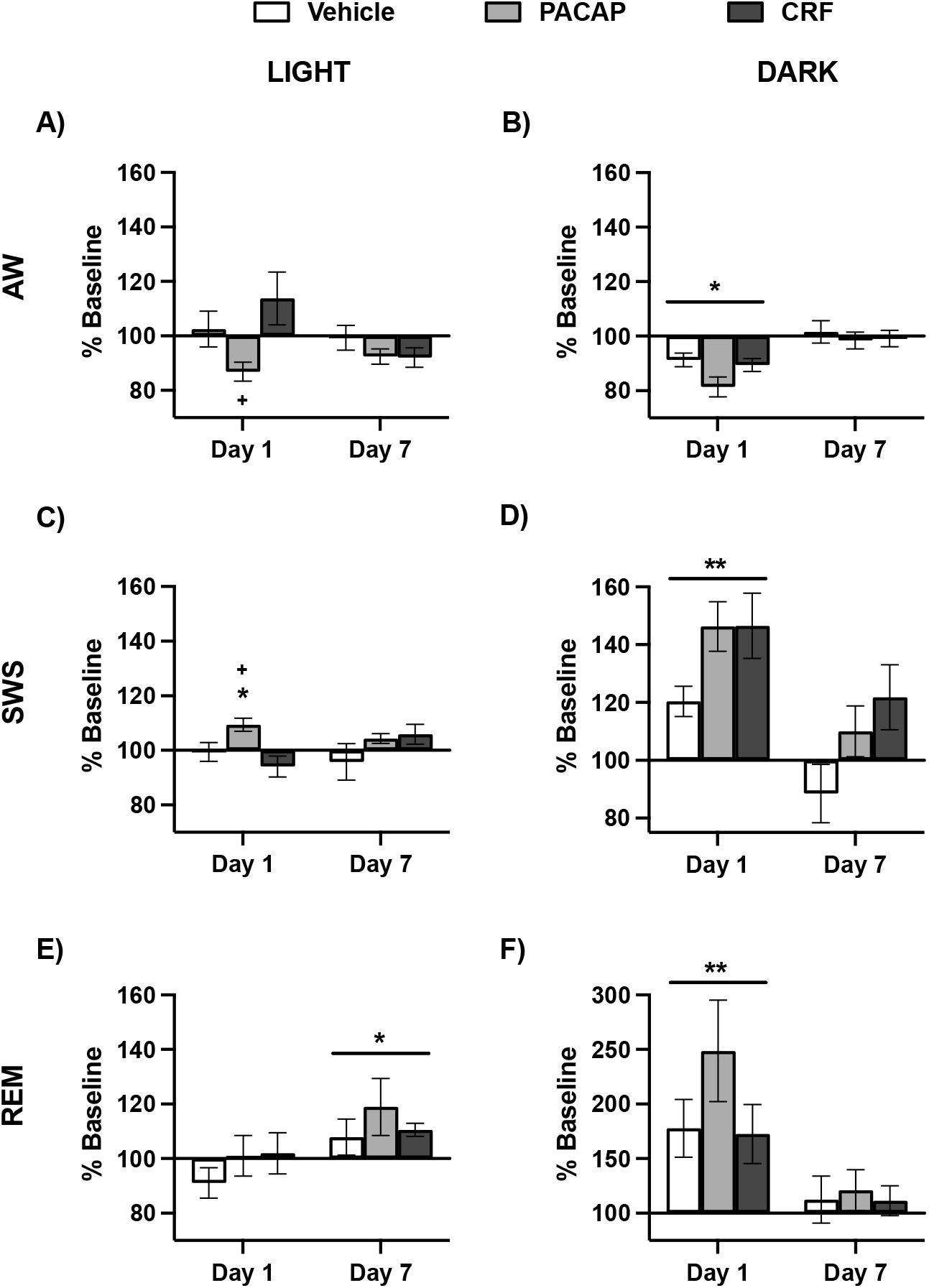
Light/Dark phase changes in vigilance states. Changes as percentage of Baseline (± SEM) after Vehicle (white), PACAP (grey) and CRF (black) treatment in duration of **A)** AW during the Light phase, **B)** AW during the Dark phase, **C)** SWS during the Light phase, **D)** SWS during the Dark phase, **E)** REM during the Light phase, and **F)** REM during the Dark phase. Asterisks indicates significant differences from Baseline vigilance states, a line with asterisks indicates that timepoint significantly differs from all other timepoints, and plus signs indicate significant differences compared to Vehicle-treated animals. * ^+^*P*< 0.05, ** *P*< 0.01.

**Figure 4:**
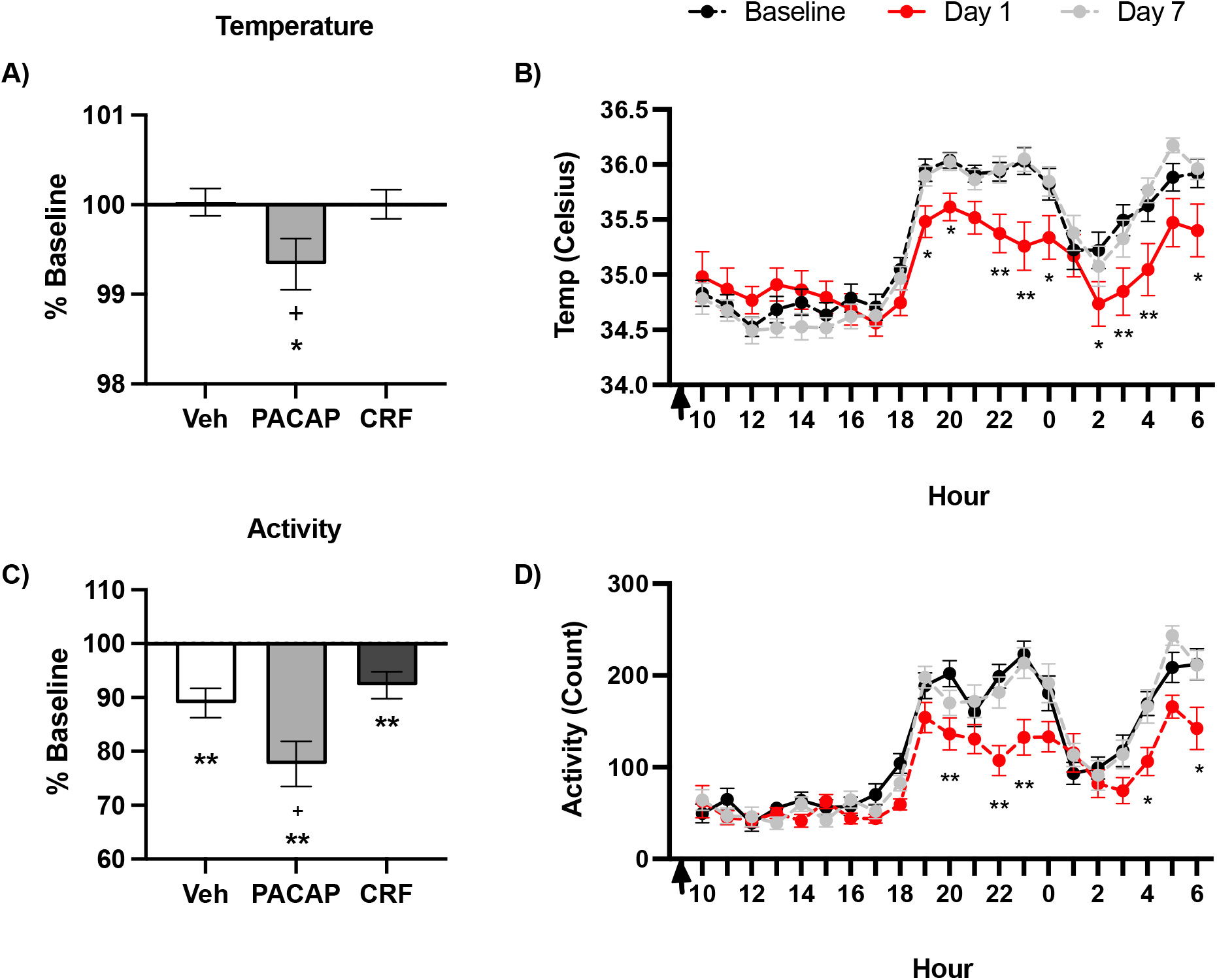
Changes in temperature and activity. A) Average (± SEM) core body temperature at Day 1 as a percentage of Baseline. B) Hourly core body temperature (± SEM) in degrees Celsius of PACAP-treated animals at Baseline (black), Day 1 (red), and Day 7 (grey). C) Average (± SEM) activity at Day 1 as a percentage of Baseline. D) Hourly activity count (± SEM) of PACAP-treated animals at Baseline, Day 1, and Day 7. Asterisks indicates significant differences from Baseline, plus signs indicate significant differences compared to Vehicle-treated animals. * ^+^ *P* < 0.05, ** *P* < 0.01.

The pattern of changes in locomotor activity after peptide treatment resembled that of changes in body temperature, although general reductions were observed in all conditions. A One-way ANOVA revealed an effect of treatment (F(2,49)=5.40, *P*=0.0076). Post hoc comparisons (Tukey’s) revealed that PACAP-treated mice differed significantly from both Vehicle (*P*=0.048) and CRF-treated animals (*P*=0.0092). T-test comparisons of Baseline activity to Day 1 activity within each treatment revealed reductions in locomotor activity levels in all groups at Day 1 compared to Baseline: Vehicle (t(16)=4.025, *P*=0.001), PACAP (t(18)=5.33, *P* < 0.0001), and CRF (t(15)=3.074, *P*=0.0077).). More detailed (1-hr bins) analysis of locomotor activity revealed a similar pattern to that observed with body temperature. In PACAP-treated animals, there were reductions in locomotor activity during the dark phase of the light cycle (**Fig. 4D**).

### EEG Power

Vigilance state durations are determined by EEG and EMG signals, but EEG signals alone can provide insight into patterns of neural activity and are known to change in response to stress in mice^10^. There is also evidence that EEG signals are dysregulated in individuals with depression^61^. We focused on frequency bands corresponding to delta (0.5-4 Hz), theta (4-8 Hz), alpha (8-12 Hz), beta (16-24 Hz), and gamma (30-80 Hz) power. Changes in EEG spectral power were calculated as a %Baseline at Day 1 for each powerband during each vigilance state.

For AW, one-way ANOVAs of each powerband revealed treatment effects on delta power (F(2, 49)=4.20, *P* = 0.021) (**Fig. 5A**). Dunnett’s post-hoc comparisons of AW delta for PACAP- and CRF-treated mice to Vehicle mice found no significant differences. T-tests comparing Day 1 to Baseline within Condition and vigilance state revealed that CRF-treated mice AW Day 1 significantly differed from Baseline for delta (t(15)=4.28, *P*=0.0007) and theta (t(15)=3.55, *P* = 0.0029) power. Alpha power at Day 1 significantly differed from Baseline in all conditions: Vehicle, (t(16)=2.57, *P*=0.021), PACAP (t(18)=2.91, *P*=0.0094), and CRF (t(15)=4.63, *P*=0.0003). AW beta power also differed from Baseline at Day 1 in Vehicle (t(16)=2.25, *P*=0.039), PACAP (t(18)=3.03, *P*=0.0072), and CRF (t(15)=2.36, *P*=0.032), as did gamma: Vehicle (t(16)=2.40, *P*=0.029, PACAP (t(18)=5.32, *P* < 0.0001), and CRF (t(15)=3.26, *P*=0.0053). Such broad changes may reflect non-specific changes in AW as a result of the ICV injection during the light phase, when mice are normally more likely to be sleeping.

**Figure 5:**
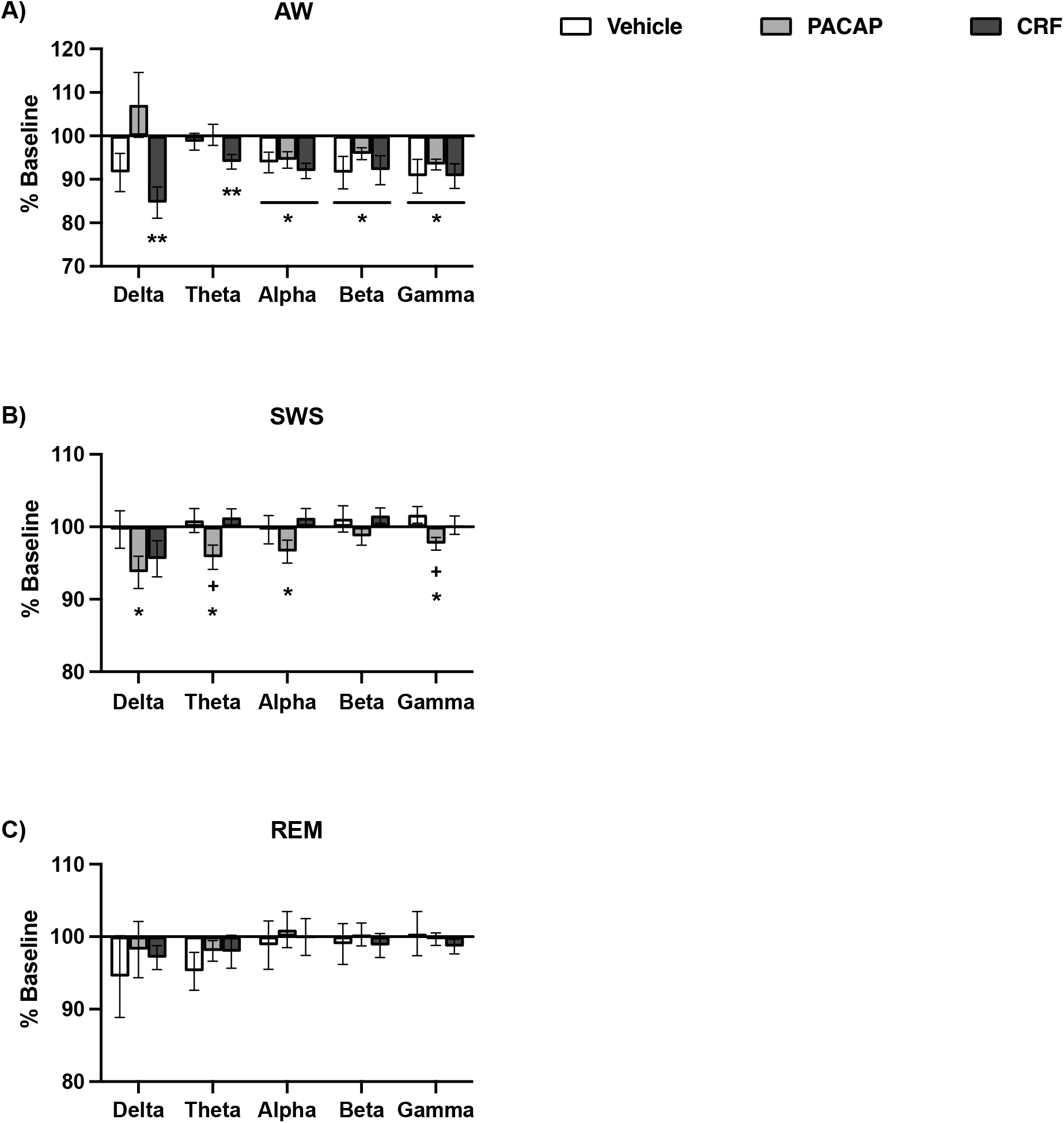
Changes in EEG absolute power by vigilance state. Average changes in EEG spectral power, displayed from low frequency (delta, 0.5-4 Hz) to high frequency (gamma, 30-80 Hz), at Day 1 as a percent of Baseline (± SEM) for **A)** AW, **B)** SWS, and **C)** REM. Asterisks indicates significant differences from Baseline, plus signs indicate significant differences compared to Vehicle-treated animals. * ^+^ *P* < 0.05, ** *P* < 0.01.

For SWS, one-way ANOVAs of each powerband revealed treatment effects on delta (F(2, 49)=4.07, *P*=0.023) and gamma power (F(2,49)=3.57, *P*=0.035) (**Fig. 5B**). Dunnett’s post-hoc tests comparing PACAP- and CRF-treated mice to Vehicle animals at these powerbands found that PACAP significantly differed from Vehicle for both theta (*P*=0.041) and gamma power (*P*=0.021). Within condition t-tests found that PACAP-treated mice significantly differed from Baseline at delta (t(18)=2.80, *P*=0.012), theta (t(18)=2.50, *P*=0.022), alpha (t(18)=2.16, *P*=0.044), and gamma powerbands (t(18)=2.67, *P*=0.016). Vehicle- and CRF-treated mice did not display changes from Baseline at any powerbands for SWS.

For REM, One-way ANOVAs and within conditions t-tests of EEG absolute power did not reveal any significant changes from Baseline at Day 1 for any condition (**Fig. 5C**).

## DISCUSSION

These studies provide a direct comparison of the effects of PACAP and CRF on sleep architecture under identical testing conditions, at dosages that cause similar anxiogenic like effects in mice. Acute treatment with PACAP broadly impacted rhythms of sleep, body temperature, and locomotor activity. CRF, conversely, did not lead to major alterations in any of these endpoints, although there were small and transient changes (e.g., SWS within the dark phase) that support the biological efficacy of the dosage tested. The changes observed in PACAP-treated mice closely resemble those previously observed in mice undergoing chronic social defeat stress (CSDS)—including decreased AW, increased SWS, increased REM sleep, and reduced amplitude of body temperature and locomotor activity rhythms^10^. Some changes were seen across all conditions, including increased locomotor activity in the 24 hours after injection, decreased wake/increased sleep during the dark phase of the light cycle and reduced alpha, beta and gamma power during AW. These broad changes might be due to the disruption light phase sleep early in the light cycle, when the ICV injection of either vehicle or peptide was administered, or the mild stress of receiving ICV injection, despite prior handling to promote acclimation. Across conditions, changes in vigilance state durations occurred primarily in the dark phase of the light cycle, with increased sleep and decreased wake when mice are typically awake and active. Within PACAP-treated animals, changes were also observed in the light phase of the light cycle, with the same pattern of decreased wake and increased sleep, contributing to the overall change in vigilance state durations observed in the PACAP condition. Corresponding with decreases in time awake, PACAP-treated animals displayed an overall decrease in temperature and locomotor activity during the 24 hours after treatment, specifically during the dark phase of the light cycle, over 9 hours after PACAP administration. The decreased peak (corresponding to amplitude) of normal circadian temperature and activity aligns with our previous work characterizing the effects of CSDS in this endpoint^10^. Changes in EEG spectral power during SWS were found in PACAP-treated animals at nearly all frequencies. Reduction and flattening of EEG power circadian rhythmicity has been associated with stress experiences in other rodent models^6,10^. Together, these findings suggest that PACAP plays an important role in stress-induced changes of sleep architecture and associated biological rhythms, whereas equipotent concentrations of CRF seem less involved in the regulation of these endpoints.

Analyses of baseline sleep prior to treatment revealed significant sex differences in duration of AW and SWS, but not REM sleep. Sex differences in sleep architecture have been reported in prior studies of both humans and rodents^7,62,63^. Extensive work indicates that sleep alterations in females are largely dependent on sex-related hormones, estrous cycle in rodents and menstrual cycle in humans^64–66^, which were not studied in these experiments. Variations in core body temperature can be utilized to determine estrous phase in female mice^56,67^. Despite access to this data via the wireless transmitters, examining the role of estrous phase on sleep was not a primary goal of this work. As such, the number of females in each estrous phase at the time of baseline was not explicitly controlled, and coincidental alignment was insufficient to properly assess a role of estrous phase on sleep/wake durations. Despite insufficient variation to analyze female sleep by estrous phase, sleep/wake durations of all females together were significantly different than males. In the future, studies could be designed to more explicitly control estrous phase, thereby enabling more detailed conclusions about how this may affect sensitivity to stress peptides.

Comparisons of sleep architecture after peptide treatment expressed as percentage of baseline did not reveal significant sex differences. This is unexpected since there are various known sex differences in PACAP and CRF systems, in addition to baseline sex differences in sleep found in this study. Sex differences have been found in CRF receptor density in stress- and anxiety-related brain regions, including the amygdala, as well as increased CRF activation after stress in females compared to males^37,39,41,42^. Expression of the cognate PACAP receptor (PAC1R) is regulated by estrogen, and has been shown to vary across the female estrous cycle in rodents^20,22^. In human clinical populations, PACAP is associated with depression in males and anxiety-related disorders in females^22,23,31^. Single nucleotide polymorphisms (SNPs) relating to PACAP and CRF receptors are associated with PTSD and symptom severity, particularly in women^22,31,68^. Furthermore, biological sex is a significant risk factor in stress- and fear-related psychiatric illnesses, with higher prevalence of PTSD, GAD, and MDD in women compared to men^35,36,69^. Despite these documented differences, we did not observe sex differences in sleep architecture as a result of PACAP or CRF under the present testing conditions. Accordingly, to best assess the overall effects of PACAP and CRF on sleep architecture, males and females were combined for analyses^58^.

Our finding that acute PACAP treatment produces increases in both SWS and REM sleep aligns with prior research on the effects of PACAP on sleep. Previous studies in rodents similarly reported that ICV and intra-pons infusions of PACAP produce increases in REM sleep^46,47^. Prior work has also shown that CSDS produces increased REM sleep, and increased REM sleep duration is characteristic of individuals with depression and a risk factor for relapse for those in currently remission^10,59^. The lingering effects of PACAP on REM sleep at Day 7, a week after PACAP administration, also aligned with our initial hypotheses. Behavioral effects of PACAP have been found to last longer than a week, and microinfusions of PACAP into the pons altered sleep for longer than a week^21,29,30,47^. We previously reported that changes in REM sleep as a result of CSDS also persist for at least five day after stress termination^10^—the longest time point tested—raising the possibility that PACAP plays a role in long-lasting changes to sleep architecture after this type of stress regimen. In addition, the paraventricular nucleus (PVN), a critical sleep region, and the parabrachial nucleus (PBn), an area associated with stress-sleep interactions, are both sites of PACAP production^27,47,56,70^. PACAP is also involved in circadian rhythms and responses to light, acting along the neuronal pathway from the retina to the superchiasmatic nucleus (SCN)^71^, representing another mechanism by which it may contribute to sleep architecture. Considered together, these findings comprise accumulating evidence to suggest that PACAP plays a key role in the acute and persistent effects of stress on sleep.

Surprisingly, we did not observe an effect of acute CRF treatment on sleep architecture, despite previous reports and evidence that CRF interacts with neural circuits involved in sleep^43–45,50,72^. While many methodological variations could account for these differential findings, the dosage of CRF and time of day of CRF administration may be involved. In the current studies, peptide dosages were selected on the basis of their ability to cause and equivalent behavioral response in the EPM, a widely-used and thoroughly-validated procedure for assessing anxiety-like behavior^57^. Region-specific infusions of CRF into the central amygdala of rats reportedly reduces REM sleep duration when provided at a very low dose, but interestingly, not at higher doses^50^, suggesting an inverted U-shaped function. While our injections were not region specific, it is possible that the dosage we used was too high, despite causing a behavioral response equivalent to that seen with PACAP. In fear conditioning studies, ICV administration of CRF has been found to enhance fear-related reductions in REM sleep in the dark phase of the light cycle, but did not alter light phase sleep immediately after treatment^43^. One of our primary objectives for these studies was to examine the persistence of PACAP and CRF effects, considering differences in other behavioral endpoints. However, the effects of PACAP on sleep architecture were largely observed in the dark phase of the light cycle, as were the only observed CRF-induced changes. The time of peptide administration in this study, early in the light cycle, could interfere with the dark phase effects of CRF. While CRF produced a dark-phase change in SWS, it is conceivable that administration of CRF later in the light phase of the light cycle, or even in the dark phase, may elicit greater alterations than we observed in the present studies. In humans, intravenous administration of CRF decreases SWS, increases wakefulness, and increases REM, particularly in women^73,74^. Existing preclinical work to elucidate the role of CRF on the effects of stress on sleep have successfully blocked stress effects with CRF antagonists, yet promising preclinical studies on this class of drugs for stress-related disorders have not led to corresponding success in humans^25,75^. The unsuccessful development of CRF antagonism monotherapy suggests that additional targets—such as PACAP systems—may be involved in the full scope of stress-sleep interactions.

Consistent with the existing literature, our findings suggest that PACAP contributes to the effects of stress on sleep, by reducing time awake and increasing sleep duration. Direct comparison of PACAP and CRF provides clear evidence that PACAP administration produces acute alterations in sleep within 24 hours, with persistent effects on REM sleep. While CRF may contribute to some circadian disruption, it does not have a comparable impact to PACAP. In general, endpoints including sleep, body temperature, and locomotor activity as endpoints for stress-related preclinical research have considerable translational value, as they are defined and measured in the same ways across species and are increasingly available in human studies through devices, such as smart phones and wearables^12^. Our findings raise the possibility that PACAP, but not CRF, plays a role in the persistence of stress-induced sleep dysregulation, as sleep problems tend to be persistent and intractable in individuals with stress-related psychiatric conditions^8^. Improved understanding of the contributions of stress peptides to sleep problems will allow for progression of treatment and intervention for sleep dysregulation in psychiatric disorders.

## ACKNOWLEDGEMENTS AND DISCLOSURES

This research was supported by P50MH115874 (to WC) and a Phyllis and Jerome Lyle Rappaport Foundation Mental Health Fellowship (to AF). Within the past 2 years, WC has served as a consultant for Psy Therapeutics. Currently, GM is an employee of Cerevel Therapeutics and KM is an employee of Jazz Pharmaceuticals, but their contributions to this work occurred while employed at McLean Hospital. None of the other authors report biomedical financial interests or potential conflicts of interest.

